# Phenological characterization of Mozambican native caprine breeds assessed through morpho-biometric traits and zoometric indices

**DOI:** 10.1101/2025.08.15.670476

**Authors:** Deiby T. Culhe, Matilde F. Matola, Élio J. R. Muatareque, Milton A. Morrombe, Matilde A. Manhique, Ramos J.Tséu, Abílio P. Changule, Maria da G. Taela, Custódio G. Bila, Manuel Garcia-Herreros

**Affiliations:** Department of Animal Production and Food Technology, Faculty of Veterinary Medicine, Eduardo Mondlane University (UEM), Maputo 1304, Mozambique; Center for Genetic Resources and Animal Assisted Techniques (CRGTRA), Directorate of Animal Science (DCA), Agricultural Research Institute of Mozambique (IIAM), Matola 1410, Mozambique; Chobela Research Station, Centro Zonal Sul, Agricultural Research Institute of Mozambique (IIAM), Magude 1121, Mozambique; Department of Animal and Public Health, Faculty of Veterinary Medicine, Eduardo Mondlane University (UEM), Maputo 1304, Mozambique; Department of Research and Development, Intermed Mozambique Lda, Maputo 1304, Mozambique; Center of Excelence in Agri-Food Systems and Nutrition (CEAFSN) - Eduardo Mondlane University (UEM), Maputo 257, Mozambique; Faculty of Veterinary Medicine and Animal Science, Save University (UniSave), Gaza Delegation, Chongoene 1206, Mozambique; National Institute for Agricultural and Veterinary Research (INIAV), Santarém 2005- 424, Portugal; CIISA-AL4AnimalS, Faculty of Veterinary Medicine, University of Lisbon, 1300-477 Lisbon, Portugal

**Keywords:** Goats, Morphometry, Native breeds, Zoometry, Mozambique

## Abstract

This study aimed to characterize the morpho-structural traits of indigenous goats reared at the Chobela Research Station in the Magude District of southern Mozambique. A total of 135 goats were randomly selected, comprising 77 Landim and 60 Pafúri animals. Racial characteristics were assessed through visual inspection, while morphometric traits were measured using a zoometric tape. Descriptive statistics and independent samples *t*-tests were performed at a 5% significance level using SPSS version 27. In terms of racial traits, all Pafúri goats exhibited a convex facial profile, whereas Landim goats showed both convex (57.9%) and concave (42.1%) profiles. Approximately 75% of the goats presented a uniform coat colour. Morphometric comparisons revealed that Landim goats had higher average values for horn length (mean difference, MD: 0.21 cm), withers height (MD: 0.14 cm), and body length (MD: 0.24 cm). In contrast, Pafúri goats had greater head length (MD: 0.18 cm), head width (MD: 0.11 cm), ear length (MD: 0.09 cm), and thoracic perimeter (MD: 0.26 cm). Regarding zoometric indices, Landim goats recorded higher body (1.71), cephalic (0.31), and thoracic (1.31) index values, while Pafúri goats exhibited a slightly higher proportionality index (0.40). The morpho-structural differences identified between Landim and Pafúri goats demonstrate distinct phenotypic profiles that can support breed characterization. These results provide a valuable baseline for future genetic studies and contribute to conservation and sustainable utilization strategies for indigenous goat populations in Mozambique.

## Introduction

Goats (*Capra hircus*) are among the earliest domesticated livestock species and have supported human livelihoods for over 10,000 years (Zeder, 2017). Their remarkable adaptability enables them to thrive across a wide range of agroecological zones, from arid deserts to tropical forests and highlands (FAO, 2021). Globally, more than 1,000 recognized breeds contribute to diverse production systems, offering meat, milk, fiber, skins, and fulfilling socio-cultural roles (Galal, 2020; Haile et al., 2022).

In many developing countries, particularly in sub-Saharan Africa, goats are a cornerstone of smallholder farming systems. They enhance food security, support income diversification, and serve critical functions in traditional ceremonies, dowries, and as a form of insurance against crop failure (Peacock, 2005; Alemayehu et al., 2022). In Mozambique, the goat population has grown steadily over the past decade and is largely composed of indigenous breeds raised under extensive or semi-extensive management systems, characterized by minimal external inputs and limited selective breeding (Mataveia et al., 2019; FAOSTAT, 2023).

Despite their socio-economic and cultural significance, the genetic and phenotypic diversity of Mozambique’s indigenous goat populations remains insufficiently documented. Phenotypic characterization, particularly of morphometric and breed-specific traits, is a critical step in the identification, conservation, and improvement of local animal genetic resources (FAO, 2012; Gizaw et al., 2020). Such efforts are fundamental to the development of sustainable breeding programs and the prevention of genetic erosion.

Considering the limited phenotypic data available, this study aimed to characterize the morpho-structural traits of two indigenous goat populations Landim and Pafúri reared at the Chobela Zootechnical Station in Magude District, southern Mozambique. It was hypothesized that, despite coexisting in the same environment, the two populations would exhibit distinct morpho-structural profiles reflecting their different genetic backgrounds and adaptive histories.

## Materials and methods

### Ethical statement

This research was performed in strict accordance with the recommendations in the legal framework for all Mozambican Public and Private Laboratories and Higher Education Institutions. The study was conducted according to the guidelines of the Declaration of Helsinki and following the Code of Ethics for animal experiments as reflected in the ARRIVE guidelines available at https://arriveguidelines.org/arrive-guidelines/study-design/1a/explanation (accessed on 12 August 2025). This study was approved by the Bioethics Committee for the use of experimental animals at the Eduardo Mondlane, Mozambique (approval date: 20 March 2023, Code Number: CBUAE-112-UEM-MZ).

### Study location and experimental material

This study was conducted at the Chobela Zootechnical Station (CZC) (Figure 1), located in Chobela village, Magude District, southern Mozambique. The district lies between latitudes 24°59′20″ S and longitudes 32°45′10″ E, and is bordered by the districts of Chókwè and Bilene- Macia to the north, Moamba to the south, Manhiça to the east, and the Republic of South Africa to the west (Miambo et al., 2019).

**Figure 1:**
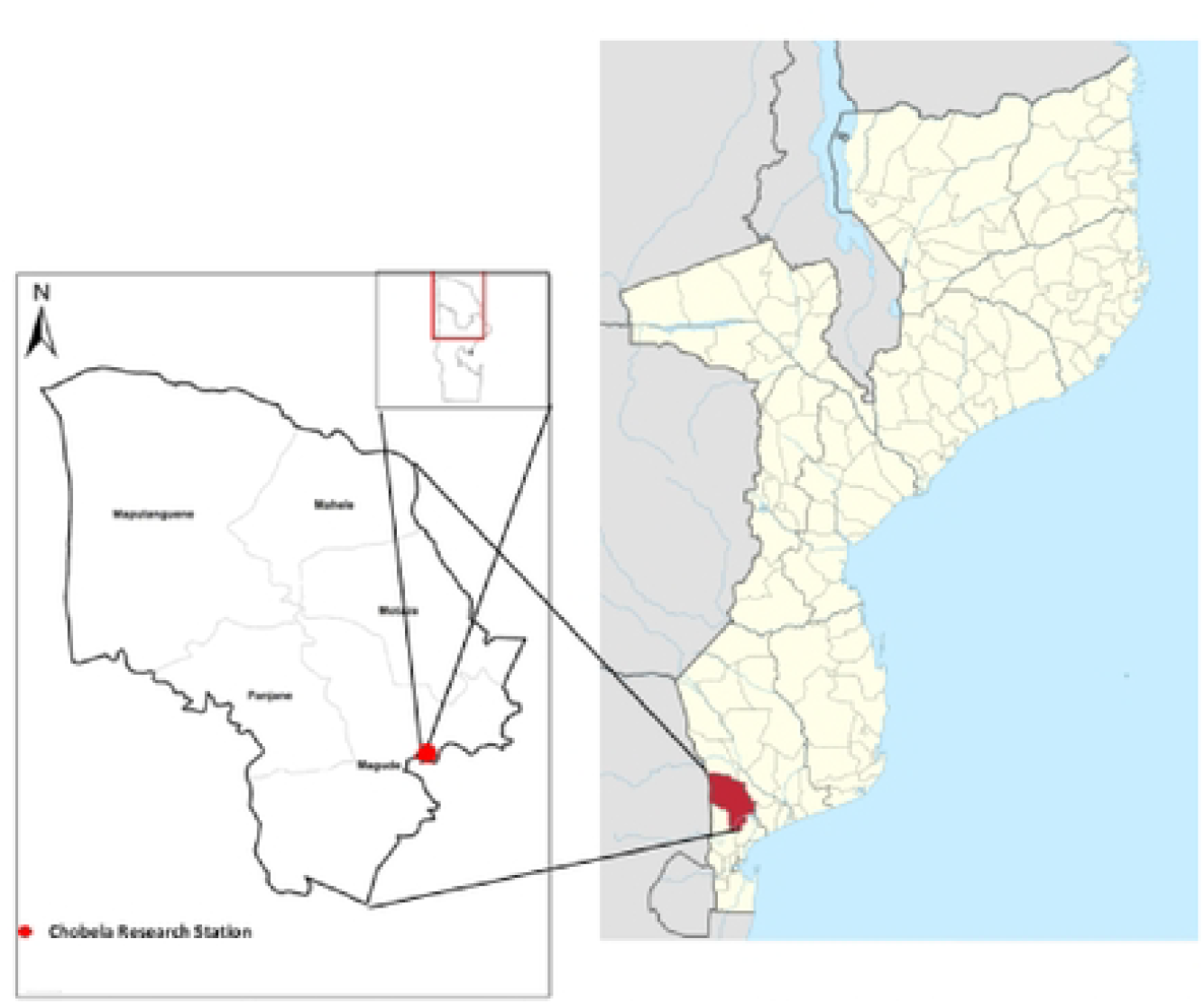
Map showing Mozambique (right) and the Chobela Research Station within the Magude District where the study was conducted.

The area supports both private and communal livestock systems, with the main species being cattle, goats, pigs, sheep, and poultry (primarily indigenous chickens). The climate is classified as dry subtropical (Köppen classification), with average annual temperatures ranging from 22 to 24°C and an average annual rainfall of approximately 630 mm. There are two distinct seasons: a hot and rainy season from October to March, which accounts for around 80% of the annual precipitation, and a cooler, dry season from April to September.

The goats were maintained under an extensive production system using natural pasture feeding. For the study, a total of 137 breeding goats were randomly selected, comprising two indigenous populations: 77 Landim (**Figure 2A:** Female goat. **2B:** Male goat) and 60 Pafúri (**Figure 3A:** Female goat. **2B:** Male goat).

**Figure 2:**
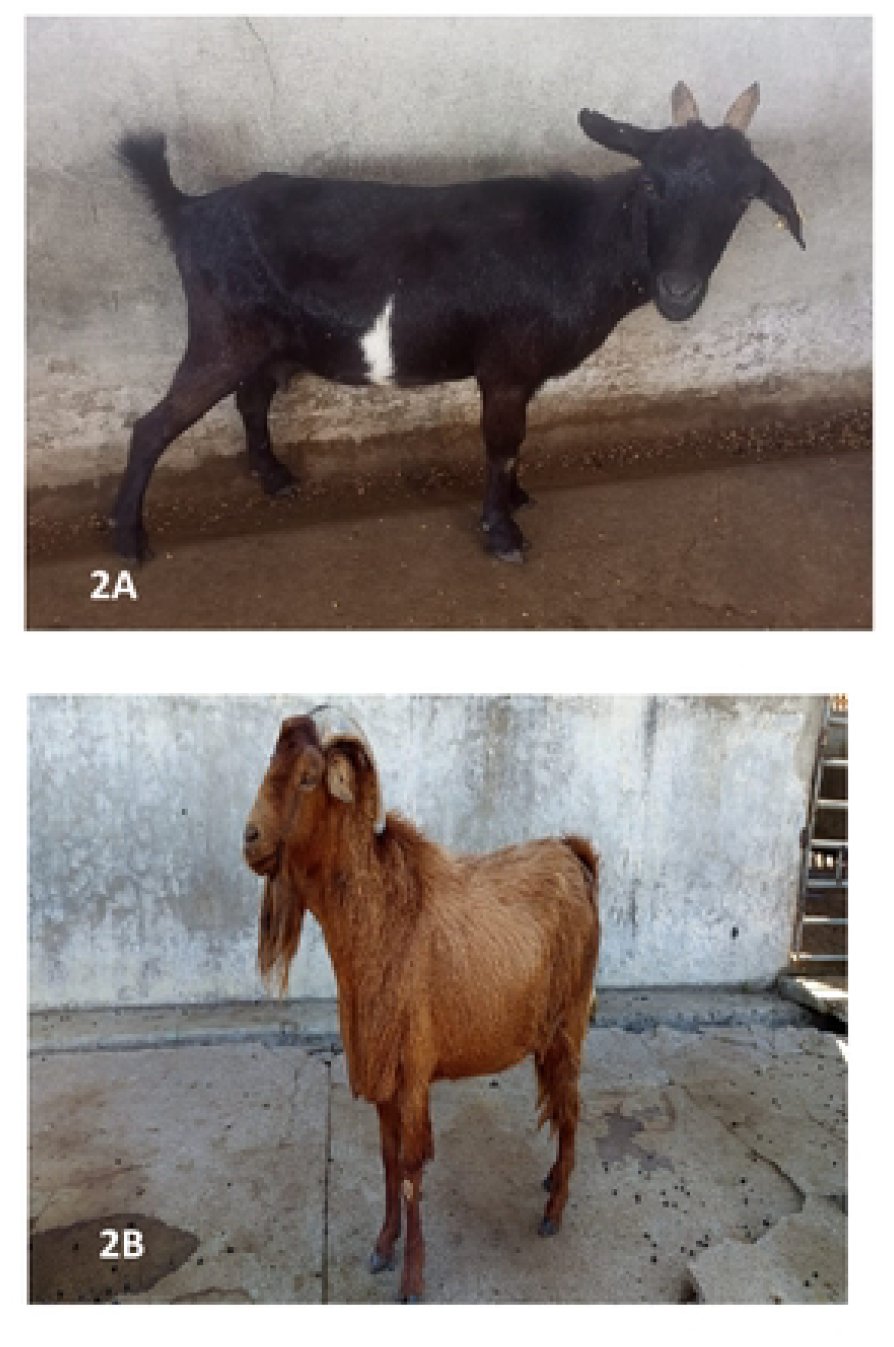
Landim goats.

**Figure 3:**
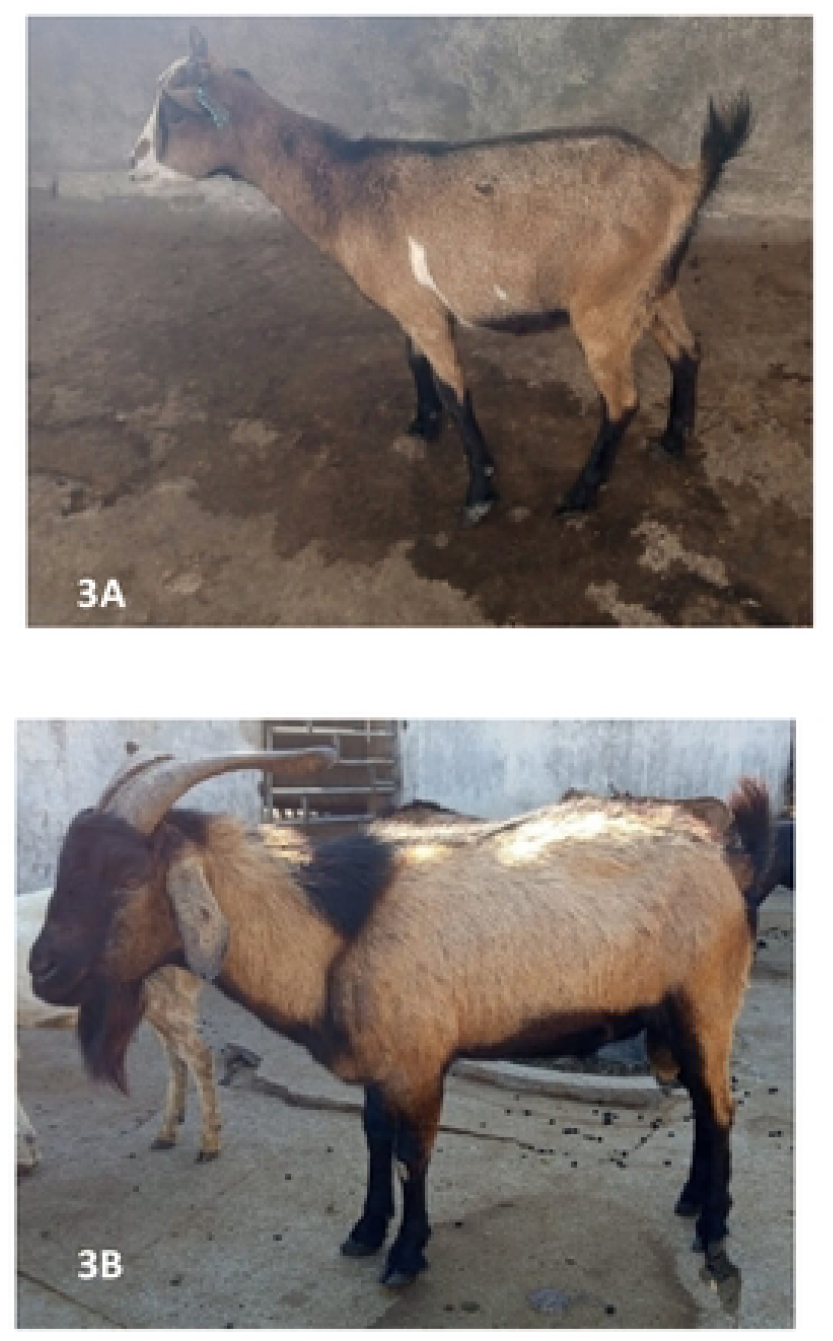
Pafuri goats.

These two indigenous breeds have been bred at the Experimental Research Station for more than 50 years. Landim goats have short-eared heads that are concave in females and slightly convex in males; horns curve backward in both sexes but are heavier in males. All males and 12% of females carry beards. Coats vary in colour (black, white, brown, or spotted), ears are erect, and the hair is short and fine. Bucks sport a short, dense mane along the backline and reach about 45 kg at two years, while does average 35 kg (Morgado, 2002; Garrine et al., 2010; Mataveia et al., 2018; Cambula and Taela, 2020). Pafúri goats display a convex head profile with divergent, well-developed horns in males and smaller, scimitar-shaped horns in females. Ears are medium-length, sometimes trimmed, with rounded tips. Both sexes have beards, a strong, well-set neck, straight back, and well-muscled limbs. Adult males weigh roughly 60 kg and females about 43 kg (Morgado, 2002; Garrine et al., 2010; Mataveia et al., 2018; Cambula and Taela, 2020).

The sample included 131 females (74 Landim and 57 Pafúri) and 6 males (3 Landim and 3 Pafúri). Animals were selected using a simple random sampling method from the breeding herds maintained at the Chobela Zootechnical Station, ensuring representation of both populations.

### Data collection

Data collection involved both visual assessment and direct physical measurements. Phenotypic characterization included the recording of racial characteristics and linear body measurements (FAO, 2012).

### Racial characterization

Morphometric traits were measured using a standard zoometric measuring tape, following proper physical restraint of each animal to ensure both data accuracy and animal and handler safety. Restraint was carried out gently, typically with the aid of halters or hand-held by trained personnel, to minimize movement and stress during measurement. This procedure aligns with the guidelines recommended by the FAO for the phenotypic characterization of animal genetic resources (FAO, 2012; FAO, 2021). The following linear body measurements were recorded (all in centimeters): Head length, head width, ear length, horn length, withers height, thoracic perimeter and body length.

### Zoometric indices

Zoometric indices were calculated using standard formulas to evaluate somatic development and infer functional conformation (Table 1). The following indices were computed as described by Salako (2006), Pires et al. (2011), Yakubu (2011), (Parés and Casanova, 2015) and Rojas- Espinoza et al. (2023).

**Table 1:**
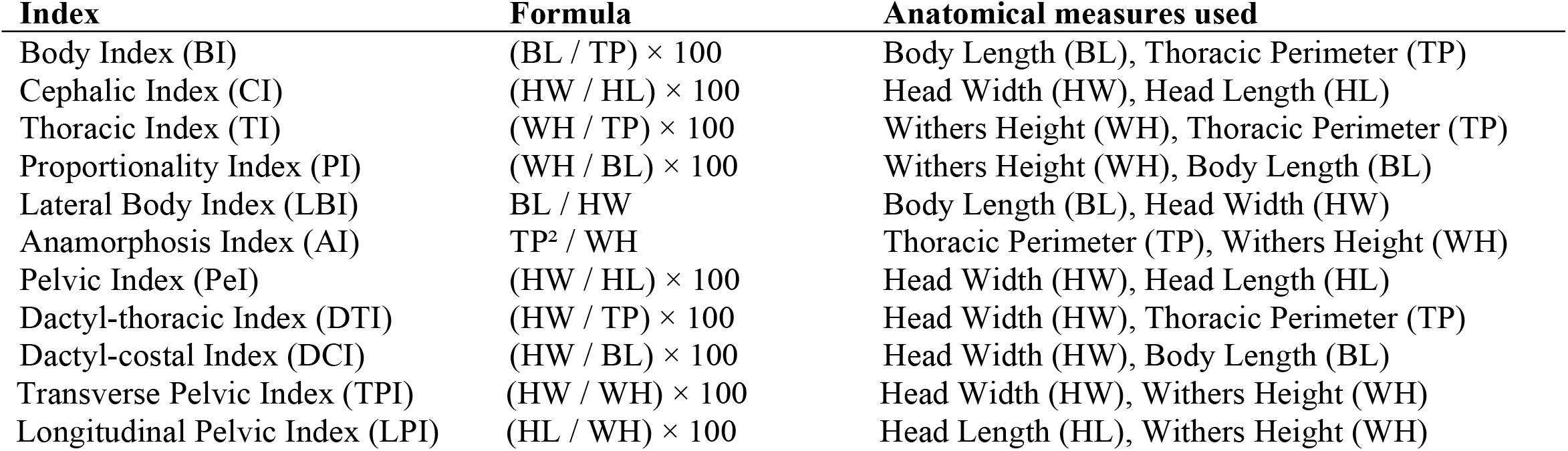
summary of calculated zoometric indices with formula.

**Table 2:** estimated frequencies and percentages of coat colour variants in Mozambican Indigenous Goat Breeds (Landim and Pafúri)

| Coat colour | Landim | Percentage | Pafuri | Percentage | Total | Percentage |
| --- | --- | --- | --- | --- | --- | --- |
| Black | 18 | 24.0% | 12 | 20.0% | 30 | 22.2% |
| Brown (light/dark) | 21 | 28.0% | 18 | 30.0% | 39 | 28.9% |
| White | 12 | 16.0% | 10 | 16.7% | 22 | 16.3% |
| Spotted/Mixed | 24 | 32.0% | 20 | 33.3% | 44 | 32.6% |
| Total | 75 | 100% | 60 | 100% | 135 | 100% |

Distribution of coat colours among Landim and Pafúri goat populations. The data represent frequencies and percentages of animals coats, based on visual assessment.

These indices were used to classify the animals according to their body conformation brevilinear, mesolinear, or longilinear and to infer their potential for meat production, particularly regarding compactness, proportionality, and skeletal development.

### Data analysis

The database was systematized in a Microsoft Excel sheet (Microsoft Corporation, Redmond, WA, USA). Subsequently, the data were subjected to descriptive statistical analyses to determine the measures of central tendency and dispersion for the respective analysis. The different data from the two breeds and sex of individuals (male and female) were subjected to assumptions of normality and homoscedasticity and then analyzed by one-way ANOVA in order to identify the potential existence of statistical significance. The ANOVA model included the interaction between breed and sex (*y =* breed + sex + breed*sex *+ e*). Tukey’s test was used to compare means. Finally, Pearson’s correlation was determined to compare pairs of morphometric parameters. All analyses were performed using the R Studio statistical software. The significance level was set at *p* ≤ 0.05).

## Results

### Racial trait characterization

Table 3 presents the morphological characteristics observed in the Landim and Pafúri goat populations, focusing on distinct facial profiles, ear orientations, horn traits, beard presence, and coat patterns.

**Table 3:**
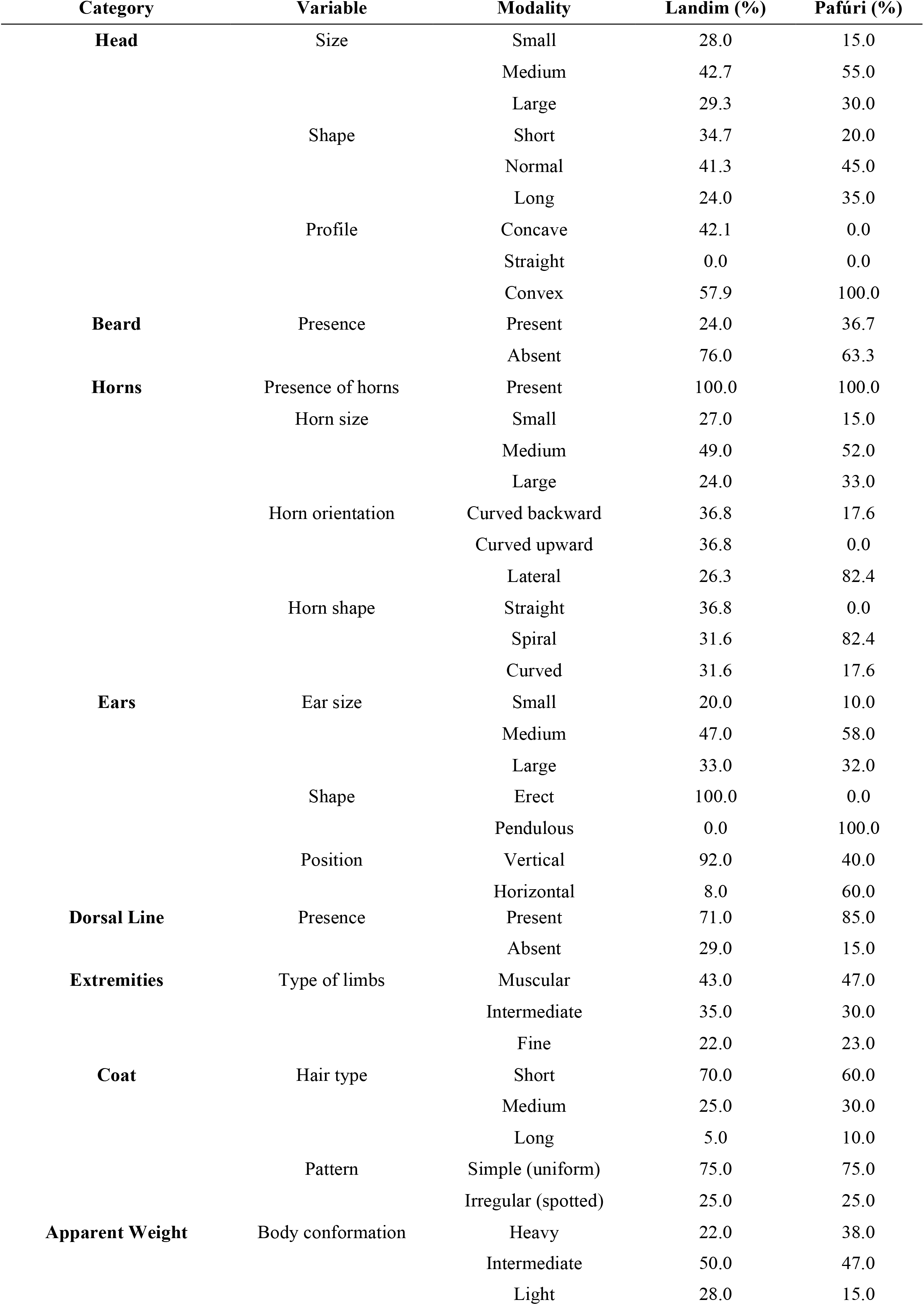
morphological characteristics observed in the Landim and Pafúri goat populations.

Regarding facial profile, the Landim goats exhibited two distinct types: convex in 57.9% and concave in 42.1% of the animals. In contrast, all Pafúri goats displayed a convex facial profile. Ear orientation differed markedly between the two populations: Landim goats had erect ears, whereas Pafúri goats presented pendulous ears.

All goats were horned, although horn length tended to be greater in the Pafúri breed. Among Landim goats, horn orientation was distributed as follows: 36.84% curved backward, 36.84% obliquely upward, and 26.3% laterally. In the Pafúri goats, 82.35% had laterally oriented horns, while 17.64% curved backward. Horn shape also varied between breeds. In the Landim population, 36.84% of animals had straight horns, while spiral and curved shapes were each observed in 31.57% of individuals. In contrast, the Pafúri goats showed a predominance of spiral horns (82.35%) and a minority with curved horns (17.64%). Beard presence was recorded in 55% of the animals, including 18 Landim and 22 Pafúri goats.

Coat pattern analysis revealed that 75% of the goats had simple patterns (uniform color), whereas 25% exhibited irregular patterns (e.g., spotting or white markings). Coat color varied widely across individuals and included solid black, light and dark brown, white, and various mottled combinations.

### Morphometric characterization

Seven linear body measurements were recorded in 131 does (74 Landim and 57 Pafúri). No statistically significant differences were observed between the two breeds for any of the morphometric traits (*p* > 0.05). The mean values (± standard deviation) for each measurement in both populations are presented in table 4.

**Table 4.** Racial characteristics observed in Landim and Pafúri goat populations.

| Trait | Landim (cm) |  | Pafúri (cm) |  | *p*-value |
| --- | --- | --- | --- | --- | --- |
|  | Male | Female | Male | Female |  |
| Head length | 17.30 ± 0.70 | 17.25 ± 0.65 | 17.44 ± 0.65 | 17.43 ± 0.75 | 0.856 |
| Head width | 11.11 ± 0.67 | 11.06 ± 0.57 | 11.27 ± 0.59 | 11.17 ± 0.59 | 0.894 |
| Ear length | 13.10 ± 0.60 | 13.08 ± 0.52 | 13.27 ± 0.59 | 13.17 ± 0.49 | 0.900 |
| Horn length | 19.22 ± 0.86 | 19.21 ± 0.83 | 19.10 ± 0.84 | 19.00 ± 0.80 | 0.856 |
| Withers height | 58.68 ± 1.31 | 58.67 ± 1.20 | 58.63 ± 1.76 | 58.53 ± 1.66 | 0.946 |
| Thoracic perimeter | 68.88 ± 1.35 | 66.94 ± 1.25 | 67.30 ± 2.17 | 67.20 ± 2.27 | 0.924 |
| Body length | 68.44 ± 1.74 | 67.31 ± 1.73 | 68.17 ± 6.10 | 67.07 ± 6.11 | 0.972 |

Values are presented as mean ± standard deviation. **TP**: Thoracic perimeter; **BL**: Body length; **WH**: Withers height; **HL**: Head length; **HW**: Head width; **HoL**: Horn length; **EL**: Ear length; **LI**: Length index; **TDI**: Thoracic development index; **TLI**: Thoraco-length index. **LI** = BL / WH × 100; **TDI** = TP / WH × 100; **TLI** = TP / BL × 100. Comparisons between breeds were tested using a two-sample t-test. Means with different superscript letters indicate statistically significant differences at $p < 0.05$ .

### Zoometric indices

Zoometric indices were calculated for all 131 does (74 Landim and 57 Pafúri). Although no statistically significant differences were observed between the two breeds (*p* > 0.05), Landim goats exhibited slightly higher mean values for the body, cephalic, and thoracic indices, suggesting a more compact and robust conformation. In contrast, Pafúri goats showed marginally higher values for the proportionality index, indicating a relatively more elongated body structure. The detailed mean values (± standard deviation) for each index are presented in table 5.

**Table 5:** Linear morphometric traits in Landim and Pafúri goats.

| Zoometric Index | Landim |  | Pafúri |  | p-value |
| --- | --- | --- | --- | --- | --- |
|  | Male | Female | Male | Female |  |
| Body Index | 99.89± 2.28 | 99.45 ± 3.16 | 99.76 ± 9.80 | 99.74 ± 9.73 | 0.829 |
| Cephalic Index | 64.60 ± 3.59 | 64.10 ± 4.09 | 63.90 ± 5.28 | 64.10 ± 4.37 | 1.000 |
| Thoracic Index | 114.10 ± 5.62 | 114.10 ± 4.72 | 115.01 ± 5.29 | 114.81 ± 5.39 | 0.532 |
| Proportionality Index | 89.68 ± 1.86 | 87.68 ± 2.86 | 88.15 ± 8.34 | 87.22 ± 8.33 | 0.691 |
| Lateral Body Index | 6.09 ± 0.45 | 6.09 ± 0.37 | 6.09 ± 0.41 | 5.99 ± 0.34 | 0.174 |
| Anamorphosis Index | 77.43 ± 3.88 | 76.40 ± 4.28 | 78.45 ± 5.60 | 78.28 ± 5.56 | 0.108 |
| Pelvic Index | 63.12 ± 4.21 | 64.12 ± 3.89 | 64.06 ± 4.20 | 64.06 ± 4.18 | 0.948 |
| Dactyl-thoracic Index | 17.49 ± 1.17 | 16.52 ± 1.11 | 16.51 ± 1.12 | 16.41 ± 1.09 | 0.606 |
| Dactyl-costal Index | 16.45 ± 1.20 | 16.43 ± 1.10 | 16.26 ± 1.14 | 16.24 ± 1.04 | 0.392 |
| Transverse Pelvic Index | 18.90 ± 1.42 | 18.85 ± 1.39 | 18.96 ± 1.54 | 18.93 ± 1.34 | 0.778 |
| Longitudinal Pelvic Index | 30.40 ± 2.30 | 29.40 ± 2.02 | 29.75 ± 2.18 | 29.70 ± 2.08 | 0.396 |

Values are presented as mean ± standard deviation. **TP: Thoracic perimeter; BL: Body length; WH: Withers height; HL: Head length; HW: Head width. Body Index = TP / BL × 100; Cephalic Index = HW / HL × 100; Thoracic Index = TP / WH × 100; Proportionality Index = WH / BL × 100; Lateral Body Index = BL / HW; Anamorphosis Index = TP<sup>2</sup> / WH; Pelvic Index = HW / HL × 100; Dactyl-thoracic Index = HW / TP × 100; Dactyl-costal Index = HW / BL × 100; Transverse Pelvic Index = HW / WH × 100; Longitudinal Pelvic Index = HL / WH × 100. Means with different superscript letters indicate statistically significant differences at $p < 0.05$ .**

Based on standard classification thresholds, Landim goats were classified as mesolinear, while Pafúri goats fell within the brevilinear category. Despite the small differences in average index values, the high variation observed in body indices and the lower variation in other indices suggest some degree of phenotypic overlap between the two populations.

## Discussion

This study was grounded in the assumption that, although raised under similar environmental and management conditions, the Landim and Pafúri goat populations would exhibit morpho-structural differences stemming from their distinct genetic backgrounds and adaptive trajectories. However, the findings revealed a high degree of similarity between the two groups, particularly in morphometric traits, for which no statistically significant differences were detected.

Nevertheless, certain trends in racial and structural attributes suggest some level of differentiation. For instance, all Pafúri goats displayed a convex facial profile, while Landim goats exhibited a broader range of horn orientations and shapes. Although these differences are subtle, they may reflect specific adaptive features or varying histories of crossbreeding. The absence of marked morphometric divergence between the populations likely indicates considerable phenotypic overlap an outcome that may be attributed to widespread uncontrolled mating and the absence of structured breeding programs. These results highlight the limitations of relying solely on phenotypic characterization to distinguish local breeds, particularly in low-input systems with minimal reproductive control.

The phenotypic variation observed in Landim and Pafúri goats reflects the richness and complexity of indigenous goat populations in Mozambique. The presence of multiple horn orientations and diverse coat patterns among Landim goats suggests substantial intra-breed variability, likely resulting from unregulated mating practices and limited genetic isolation. This finding is consistent with previous reports that attribute weak breed boundaries in many African goat populations to communal grazing systems and uncontrolled reproduction (Mekuriaw et al., 2021; Haile et al., 2022). In contrast, the uniform convex facial profile and predominance of spiral-shaped horns in Pafúri goats align with their known lineage as a crossbreed between Boer bucks and indigenous does (Cumbula and Taela, 2020). However, the lack of marked phenotypic differentiation between the two breeds in this study may indicate ongoing gene flow and the absence of structured selection programs a concern similarly raised in phenotypic assessments of indigenous goat populations across Eastern and Southern Africa (Gizaw et al., 2020; Alemayehu et al., 2022).

The absence of statistically significant differences in linear body measurements between the two breeds suggests **a** high degree of morphological overlap. Nevertheless, trends in trait dominance such as greater horn length and withers height in Landim goats, and broader heads and larger thoracic perimeters in Pafúri goats may reflect adaptive features shaped by ecological niches and local selection pressures (Galal et al., 2020; Tadesse et al., 2022).

This morphometric stability in adult animals is expected, as body measurements tend to plateau after reaching maturity, reflecting the combined effects of genetic potential and environmental adaptation (Fajemilehin and Salako, 2008). Still, even subtle differences in specific traits may hold functional or adaptive significance in local production systems and should be integrated into community-based selection programs (Kosgey et al., 2006; Haile et al., 2019).

Zoometric indices are valuable tools for assessing animal conformation and inferring production potential. In the present study, Landim goats exhibited higher values for the body, cephalic, and thoracic indices, while Pafúri goats showed slightly higher values for the proportionality index. These findings support the classification of Landim goats as mesolinear and Pafúri goats as brevilinear, indicating moderate and compact body frames, respectively both conformations being suitable for meat production (Sanudo, 2009; Getaneh et al., 2022).

While such indices are useful for comparative assessments, the high degree of morphological similarity observed between the two populations suggests that morphometric data alone may be insufficient for robust breed differentiation. Recent studies emphasize the need to integrate morphometric, genomic, and environmental data for a comprehensive characterization and sustainable management of indigenous livestock breeds (FAO, 2021; Leroy et al., 2022).

Uncontrolled mating systems remain a major constraint for breed conservation in low-input livestock systems. In such contexts, genetic erosion is accelerated by the absence of mating control and structured breeding strategies (Mwacharo et al., 2017; Gicheha et al., 2023).

## Conclusion

This study characterized Landim and Pafúri goat ecotypes based on phenotypic traits and linear body measurements under extensive conditions at the Chobela Zootechnical Station. Both ecotypes exhibited distinct racial characteristics and were classified as mesolinear, indicating moderate body frames with slight meat aptitude. These results provide a phenotypic baseline to support conservation and improvement efforts for indigenous goat populations in Mozambique.

## Acknowledgement

The authors are grateful for the support provided by the Agricultural Research Institute of Mozambque (IIAM-CZS) and all staff involved.

## Notes

### Competing Interest Statement

The authors have declared no competing interest.

